# Diverse progenitor cells preserve salivary gland ductal architecture after radiation induced damage

**DOI:** 10.1101/295501

**Authors:** Alison J. May, Noel Cruz-Pacheco, Elaine Emmerson, Kerstin Seidel, Sara Nathan, Marcus O. Muench, Ophir Klein, Sarah M. Knox

**Affiliations:** Program in Craniofacial Biology, Department of Cell and Tissue Biology, University of California, 513 Parnassus Avenue, San Francisco, CA, 94143, USA.; Department of Pediatrics and Institute for Human Genetics, University of California, San Francisco, San Francisco 94143, USA.; The MRC Centre for Regenerative Medicine, The University of Edinburgh, Edinburgh, UK; Blood Systems Research Institute, San Francisco, CA, 94118, USA.

**Author notes:** Corresponding Author Sarah M. Knox. Lead Contact: Sarah M. Knox.

**Keywords:** radiotherapy, stem cells, regeneration, salivary gland, KRT14, KIT

## Abstract

The ductal system of the salivary gland has long been postulated to be resistant to radiation-induced damage, a common outcome incurred by head and neck cancer patients receiving radiotherapy. Yet, whether the ducts are capable of regenerating after genotoxic injury, or if damage to ductal cells induces lineage plasticity, as has been reported in other organ systems, remains unknown. Here, we show that two ductal progenitor populations marked by KRT14 and KIT exclusively maintain non-overlapping ductal compartments after radiation exposure but do so through distinct cellular mechanisms. KRT14+ progenitor cells are fast cycling cells that proliferate in response to radiation-induced damage in a sustained manner and divide asymmetrically to produce differentiated cells of the larger granulated ducts. Conversely, KIT+ cells are long lived progenitors for the intercalated ducts that undergo few cell divisions either during homeostasis or after gamma radiation, thus maintaining ductal architecture in the near absence of cell turnover. Together, these data illustrate the regenerative capacity of the salivary ducts and highlight the heterogeneity in the damage responses used by salivary progenitor cells to maintain tissue architecture.

**Summary Statement:** The salivary gland ductal network is maintained during homeostasis and after genotoxic injury by diverse progenitors that respond differentially to radiation induced damage.

## Introduction

Salivary glands are composed of a complex architecture of saliva synthesizing acini and transporting ducts capable of producing up to 1.5L of this sero-mucous liquid per day (in humans). The epithelium coordinating this secretory program is very heterogeneous in cell type and consists of 2 kinds of acinar cells (mucous and serous) and at least 3 kinds of ductal cells of increasing size (intercalated (smallest), granulated/striated and excretory (largest)). These cells are morphologically and functionally distinct, allowing the organ to alter the composition of the saliva under un-stimulated and stimulated conditions (e.g. food intake). Such cell diversity implies that maintenance of tissue during homeostasis or after injury occurs via differentiation of multipotent progenitors capable of producing multiple lineages or distinct progenitor populations that act to replace a single cell type. In support of the latter, over the last few years a number of distinct progenitor cell populations have been identified to contribute to one of the multiple cell types constituting the adult epithelium. SOX2+ cells in the acini and KRT14+ cells in the ducts give rise to mucous acinar and granulated ductal lineages, respectively (Emmerson et al. 2018; Kwak et al. 2016). Recently we identified KIT as a second progenitor population for the ductal lineage that give rise to the intercalated ducts (Emmerson et al. 2018), further supporting the requirement for multiple progenitors in SG homeostasis. However, the mechanisms by which these progenitors replace ductal cells, whether they acquire lineage plasticity to replace multiple cell types as well as their ability to regenerate the ductal system after damage is unknown.

Salivary glands originate from an invagination of the oral epithelium into a condensing mesenchyme (embryonic day (E) 11.5, 6-8 weeks in humans) and form an initial pre-acinar ‘end bud’ on what will become the secretory duct for the oral cavity. This single pre-acinar end bud undergoes rapid expansion, clefting, duct formation, lumenization and terminal differentiation to form a highly branched lobular structure capable of producing saliva by birth (Tucker 2007). The murine ductal lineage emerges from a relatively undifferentiated population of KRT5+KRT19-progenitors that differentiate to form the morphologically and functionally distinct KRT8 enriched intercalated, MUC19+ granulated/striated and KRT19+ excretory duct (Knox et al. 2010; Lombaert et al. 2013). These ducts differ vastly in size with intercalated ducts connecting the acini to the ductal system being the smallest and excretory ducts residing in the connective tissue connecting the oral cavity to this system being the largest. Furthermore, the ductal cells themselves differ in size, shape and function: intercalated ducts are composed of small cuboidal cells that passively absorb ions from the saliva whereas the large columnar cells of the striated ducts actively absorb Na+ and secrete HCO3- (Lee et al. 2012).

Salivary glands are inadvertently injured from radiation treatment delivered to patients for the elimination of head and neck tumors (~60,000 new patients per year in US (Siegel et al. 2015)). Off target radiation destroys saliva synthesizing acinar cells (Redman 2008; Sullivan et al. 2005) and results in a lifetime of dry mouth and co-morbidities (e.g., oral infections, poor wound healing, dental decay (Brown et al. 1975; Dreizen et al. 1977; Dusek et al. 1996)). We recently showed that acinar progenitors can replenish the acini immediately after radiation induced damage (Emmerson et al. 2018). Intriguingly, the ductal system seems far less perturbed after radiation treatment than the acini, with ducts marked by KRT5 being similar in phenotype to non-irradiated tissue (Knox et al. 2013). Yet whether the ducts actively regenerate after radiation and if this is mediated by KRT14+ and/or KIT+ cells is unknown.

Here we show that the salivary ductal network is maintained during homeostasis and after genotoxic injury by KRT14+ and KIT+ progenitors that differ in their regenerative behavior. Using short- and long-term lineage tracing, we demonstrate that KRT14+ and KIT+ cells are unique, non-overlapping progenitor populations for the ductal lineage that replenish specific cellular compartments during homeostasis and after damage. We also demonstrate that these cells do not acquire lineage plasticity after damage, thereby limiting repair of each ductal cell type solely to its single progenitor. Finally, we show that KRT14+ and KIT+ populations differ substantially in how they preserve tissue architecture in the face of genotoxic injury, with KRT14+ cells repopulating ductal cells through asymmetric division whereas KIT+ cells maintain ductal tissue structure through limiting cell turnover.

## Results

### KRT14 and KIT segregate during development to mark distinct ductal cells in both mice and humans

Although both KIT+ and KRT14+ cells have been demonstrated to give rise to ducts in the adult SG, whether these mark the same ductal progenitor cell is unclear. To confirm their independence, we first determined the location of KIT and KRT14 protein in murine and human adult SG. In murine SG, KRT14 was expressed by a group of 4-8 KRT5+ cells wrapping around the proximal portion of the intercalated duct as well as by smooth muscle actin (SMA+) myoepithelial cells (Figure 1, A and Figure S1, A). Although located distal to the granulated duct, KRT14+ SMA- cells did not express the granulated duct cell marker MUC19 (Figure S1, B). KIT expression was observed predominately in the intercalated ducts with lower expression by a subpopulation of cells in the granulated ducts (Figure 1, A). This segregation of proteins was in contrast to the E13-E14 developing gland where KIT and KRT14 were often co-expressed by acinar cells (Lombaert et al. 2013) although by E15 few KRT14+ cells expressed KIT (Figure 1, B), suggesting segregation of these lineages occurs before terminal differentiation.

**Figure 1.**
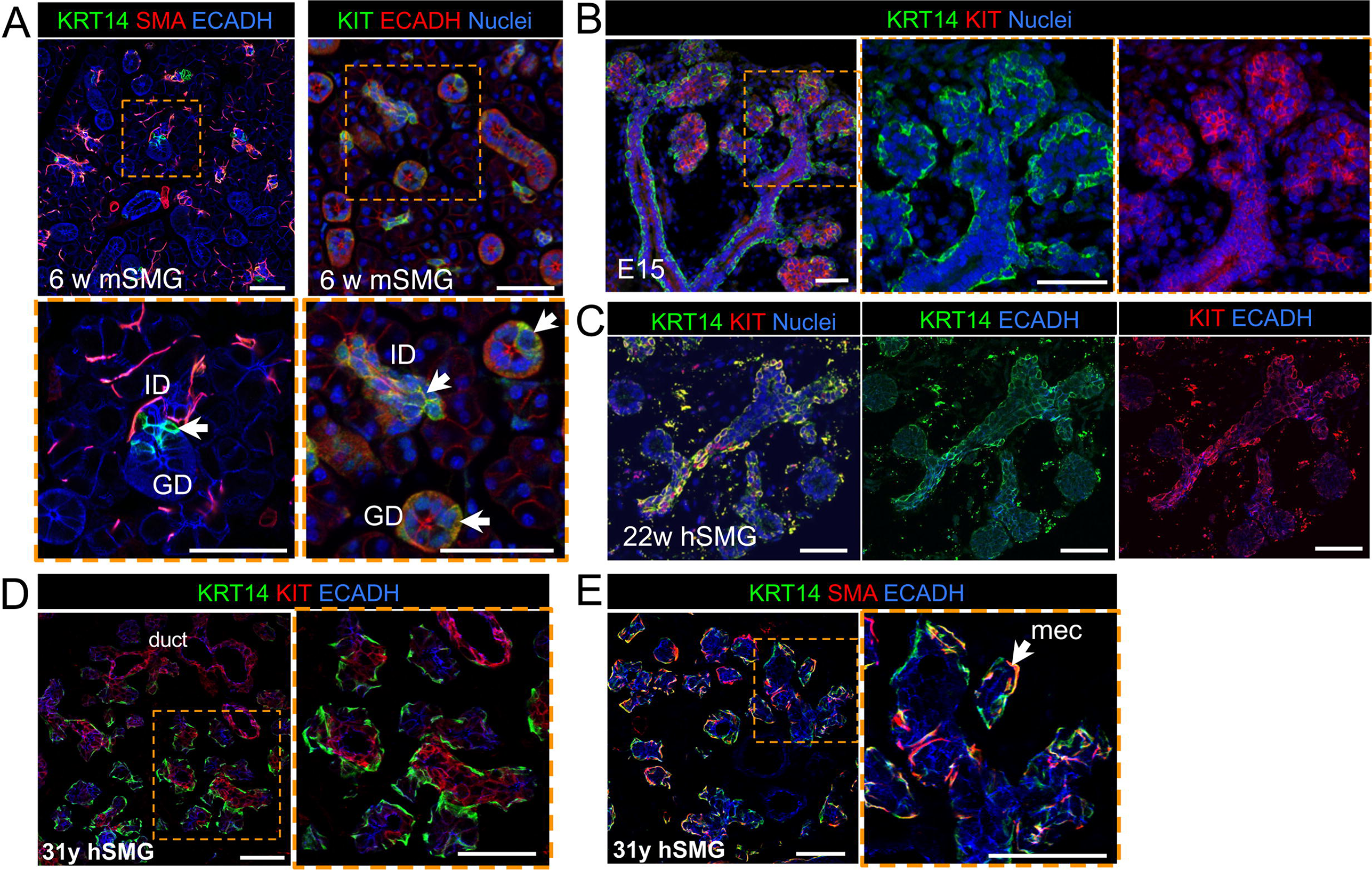
KRT14 and KIT mark distinct populations of ductal cells in murine and human SG. Murine (m) adult (A) and embryonic day 15 (E15, B) submandibular gland (SMG) or human (h) fetal (C) or adult (D and E, 31 years old) SMG immunostained for KIT, KRT14 and SMA. All scales bars are 50µm. ID = intercalated duct, GD = granulated duct, mec = myoepithelial. Arrows in (A) indicate KRT14+ SMA- ductal cells, arrows in (B) indicate KIT expression in ID and GD.

Despite KIT being a well-studied receptor in stem cell biology that has been postulated to be involved in the regeneration of salivary glands, the localization of KIT in developing or adult human SG is unclear. Similar to the murine SG, in the human SG KIT and KRT14 were co-expressed in both acinar and ductal cells during fetal development (Figure 1, C) whereas in the adult tissue KIT no longer co-localized with KRT14 and was enriched in intercalated as well as larger ductal cells (Figure 1, D-E). As for the mouse, we also observed KRT14+ cells that were either SMA+ or SMA- (Figure 1, E), indicating myoepithelial and basal duct cells, respectively. Thus, the adult mouse recapitulates the adult human system, confirming it as a useful system to model these divergent cell types.

### KRT14+ and KIT+ cells are multipotent progenitors during SG development that become unipotent at different time points to produce a single non-overlapping ductal cell type

As KIT and KRT14 co-localize during development and KRT14 is expressed by the oral epithelium, before ontogenesis, we first asked if KRT14+ cells give rise to KIT+ cells by performing genetic lineage tracing using an inducible *Krt14* promoter crossed to a *Rosa26*^*mTmG*^ reporter line. Similar to a previous study utilizing a non-inducible *Krt14* promoter (Lombaert et al. 2013), we found Cre activation at E10.5 (before invagination) and E12.5 resulted in GFP+ cells marking the entire E16.5 epithelial compartment (including acinar, duct and myoepithelial cells; Figure 2, A) and confirmed that KRT14+ progenitor cells give rise to KIT+ cells (Figure 2, A). However, when lineage tracing was initiated at post-natal stage (P) 2 (before the emergence of granulated ducts, (Srinivasan & Chang 1979)), P30 (ductal system is fully differentiated) or 6 weeks (w), KRT14+ cells contributed solely to the ductal compartment and more specifically, granulated ducts (Figure 2, B) indicating that the fate of KRT14+ cells is determined at or before P2.

**Figure 2.**
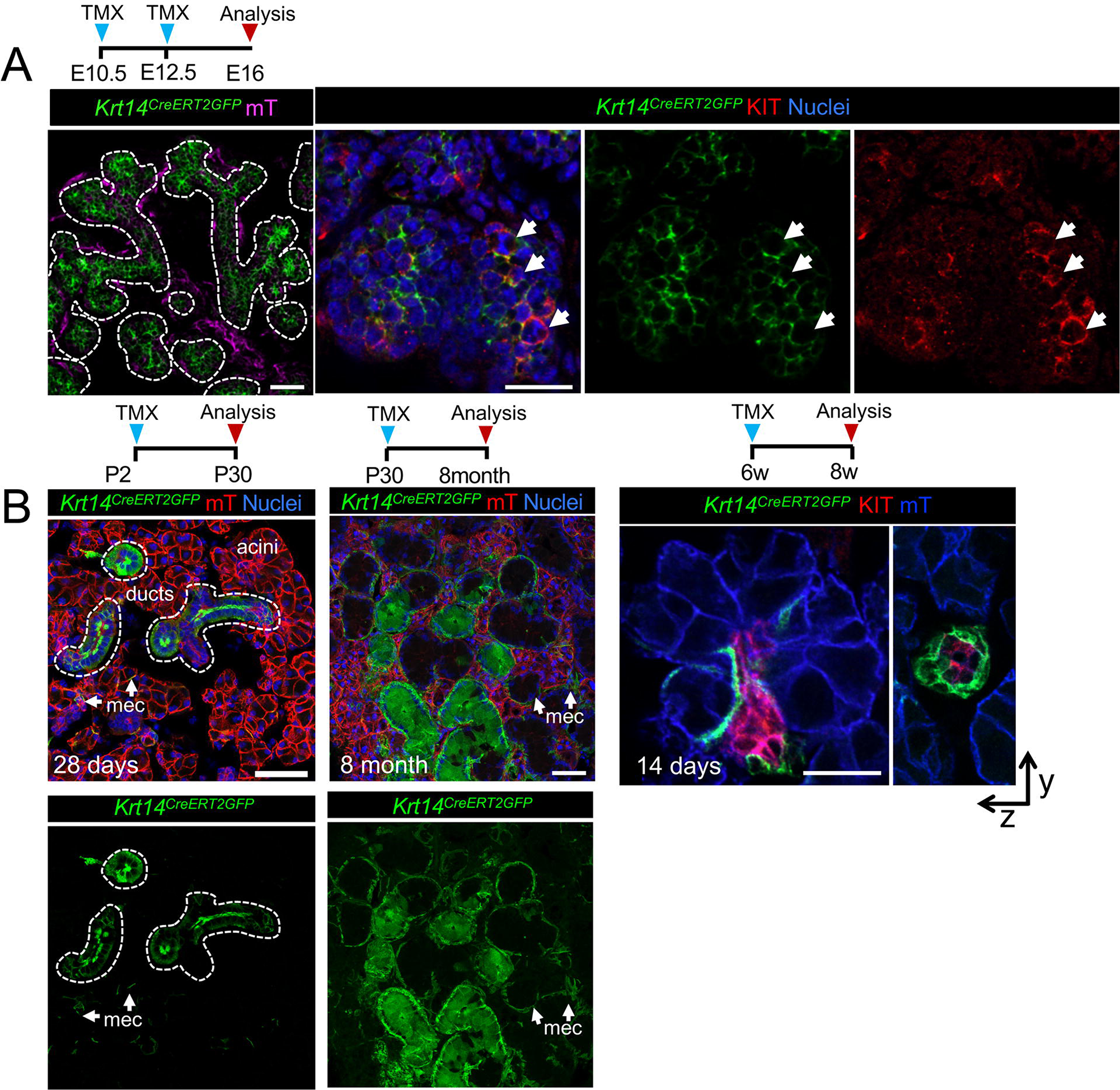
KRT14+ cells become lineage restricted to produce granulated ducts and not KIT+ intercalated ducts. Genetic lineage tracing in *Krt14^CreERT^;Rosa26^mTmG^* was activated at E10.5/E12.5 (A), or P2, P30 or 6 w (B) and cells traced for 4 days to 8 months (as indicated) before immunostaining for KIT or nuclei. All scales bars are 50µm. mec = myoepithelial cell.

Next, we tested whether KIT+ cells also became lineage restricted by P2 by utilizing the inducible *Kit* promoter (*Kit^CreERT2^*). In contrast to KRT14+ cells, activation of Cre at P2 resulted in both GFP+ acinar and duct cells (but not myoepithelial cells), indicating that KIT+ progenitors maintained their multipotency for longer (Figure 3, A). However, by 6 weeks of age, KIT+ cells replenished the intercalated ducts (KRT8+ KIT+) but not acinar cells or KRT14+ cells surrounding these ducts (Figure 3, A-C). Although we found a subpopulation of granulated duct cells were also labelled by GFP, these cells expressed endogenous KIT+ and were not GFP+ by 6 months of lineage tracing (Figure 3, C). Together, these data indicate that both KIT+ and KRT14+ cells contribute to multiple epithelial lineages of the developing SG and become lineage restricted at distinct time points to produce non-overlapping duct cell populations.

**Figure 3.**
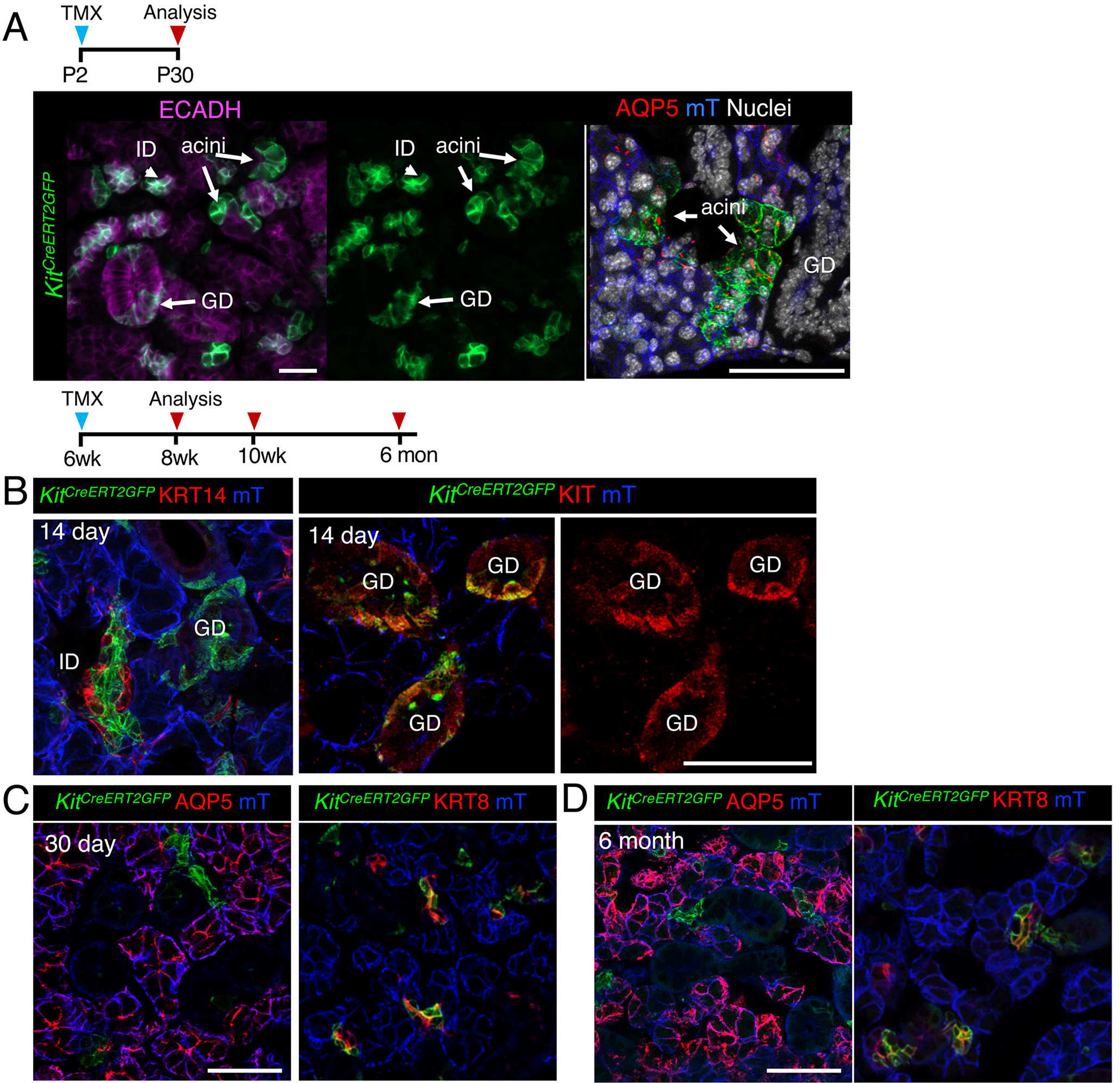
KIT+ cells in the adult SMG contribute to the intercalated ducts but not KRT14+ cells. Genetic lineage tracing in *Kit14^CreERT^;Rosa26^mTmG^* was activated at P2 (A), or 6 w (B and C) and cells traced for 14 or 30 days or 6 months (as indicated) before immunostaining for KRT14, KIT, acinar marker AQP5, duct marker KRT8 or nuclei. Scale bars are 50µm. mec = myoepithelial cell. ID = intercalated duct, GD = granulated ducts.

### KRT14+SMA+ cells give rise to myoepithelial cells and not ductal or acinar cells

As KRT14+ cells also contributed to the myoepithelial compartment (Figure 2, (Kwak et al. 2016)), we next determined if KRT14+SMA+ cells were capable of producing acinar or ductal cells by performing lineage tracing using the tamoxifen inducible *Acta2* promoter (*SMA^CreERT2^*,(Wendling et al. 2009)) crossed to an RFP reporter. During development, basal KRT14+ cells in the end bud begin to express SMA with the emergence of these cells from the acini by E16 (Figure 4, A). However, a population of KRT14+ cells within the ducts remains SMA-negative (Figure 4, A). Activation of Cre at E15 resulted in the production of SMA+ myoepithelial cells, but not KRT14+ ductal cells or AQP5+ acinar cells (Figure 4, A), suggesting that lineage restriction for the myoepithelial cell lineage occurs at a time point preceding myoepithelial emergence from the basal epithelium of the end bud. This is in contrast to the acinar lineage which we found to be derived from KRT14+ cells until E16 (Figure 4, C). To determine if SMA+ cells contributed to other epithelial lineages in adult SG, we traced cells for 30 days and 6 months but found no contribution of SMA+ cells to the ductal or acinar lineages (Figure 4, D and E), indicating that KRT14+ myoepithelial cells give rise to themselves exclusively.

**Figure 4.**
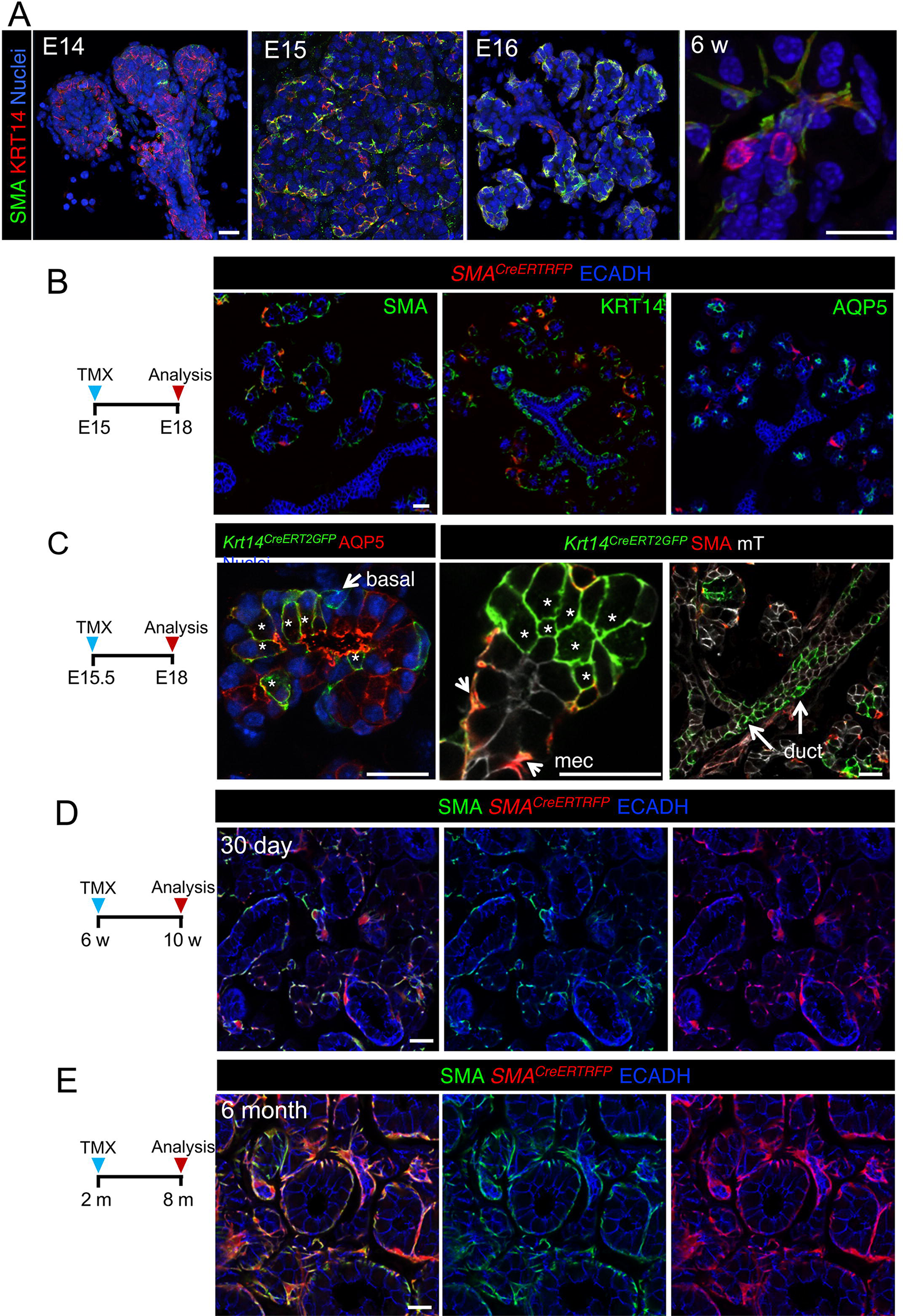
KRT14+ SMA+ cells give rise to myoepithelial cells but not duct or acinar cells. A) KRT14 and SMA localization in developing and adult SMG. B and C) Genetic lineage tracing was activated in *SMA^CreERT^;Rosa26^RFP^* and *Krt14^CreERT^;Rosa26^mTmG^* at E15 or E15.5 (as indicated). D and E) Genetic lineage tracing was activated in *SMA^CreERT^;Rosa26^RFP^* mice at 6w and traced for 6 months before immunostaining for SMA. Scale bars are 20µm. mec = myoepithelial cell, * = GFP+ acinar cell.

### KRT14+ but not KIT+ cells replenish the ductal compartment after radiation-induced damage through asymmetric division

Radiotherapy for head and neck cancer inadvertently injures the SG and causes an eventual loss of the acinar cell compartment (Knox et al. 2013; Grundmann et al. 2010). In contrast, the ductal compartment remains relatively intact suggesting that the ductal system has long term regenerative capacity (Knox et al. 2013). We tested this notion by applying gamma radiation (IR) to the neck region of mice (24h after Cre recombination) and lineage tracing for KRT14+ or KIT+ cells for 14 or 30 days. Similar to glandular homeostasis, after a single 10 Gy dose of IR KRT14+ cells gave rise to GFP+ granulated duct cells and myoepithelial cells (Figure 5, A), indicating that KRT14+ cells remain capable of repopulating the tissue but do not acquire lineage plasticity in response to genotoxic shock. Similarly, we observed GFP+ cells in the intercalated duct compartment of the irradiated *Kit^CreERT2^;Rosa26^mTmG^* SG (Figure 5, B), suggesting that they were either replenishing themselves or were capable of surviving radiation-induced damage.

**Figure 5.**
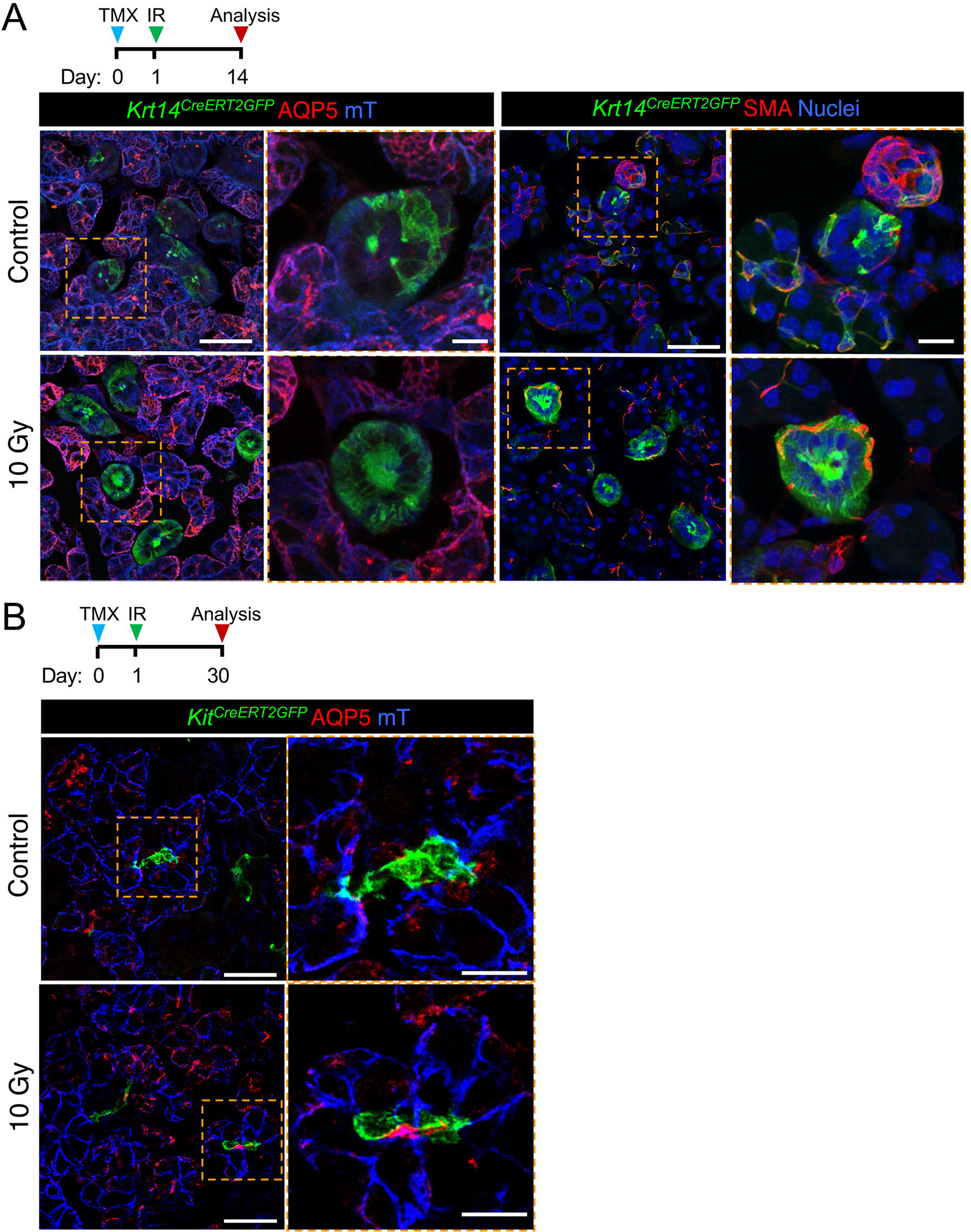
KRT14+ and KIT+ cells maintain the ductal compartment after radiation induced damage. Lineage tracing was performed in *Krt14*^*CreERT*^;*Rosa26*^*mTmG*^ (A) or *Kit*^*CreERT*^;*Rosa26*^*mTmG*^ (B) at mice at 6w with Cre activation 24 h before treatment with 0 (control) or 10 Gy of gamma radiation. SG were then traced for 14 or 30 days before immunostaining for AQP5 or SMA. Scale bars in A and B are 50 µm, orange box indicates magnified area scale bar = 20µm.

To determine if duct regeneration was mediated by proliferation of progenitor cells, we analyzed cell division of KRT14+ and KIT+ cells during homeostasis and after radiation treatment. Similar to previous studies, there was very limited cell proliferation in the ductal system of the homeostatic gland including the KIT+ intercalated ducts(Kimoto et al. 2007; Chibly et al. 2014; Kim et al. 2008) with cell division being restricted almost exclusively to KRT14+ cells (Figure 6, A). After radiation induced damage the number of dividing KRT14+ cells was initially reduced (3 days post-IR), suggestive of cell cycle exit. However, cell proliferation increased over time such that we measured a >4-fold increase in EdU+KRT14+ cells in IR SG compared to control tissue at 14 days post-IR (Figure 6, A-B), indicating that these cells undergo delayed mitosis in response to IR-induced injury. As the number of KRT14+ cells did not increase with increased proliferation (Figure 6, C) whereas GFP+ cells derived from KRT14+ progenitors increased in the granulated ducts of irradiated tissue (Figure 4, A), we conclude that KRT14+ cells repopulate the granulated ducts by asymmetric division i.e., a KRT14+ cell gives rise to itself and a differentiated daughter cell (Figure 6, D). In contrast to KRT14+ cells, we found almost no EdU+ KIT+ cells under either homeostatic or injury conditions (Figure 6, E), confirming previous reports that KIT+ cells are long lived slow dividing cells (Chibly et al. 2014). Furthermore, this outcome suggests that the intercalated duct cells are maintained rather than actively replenished after damage. Thus, KRT14+ and KIT+ cells utilize vastly different cellular mechanisms to maintain ductal architecture after injury.

**Figure 6.**
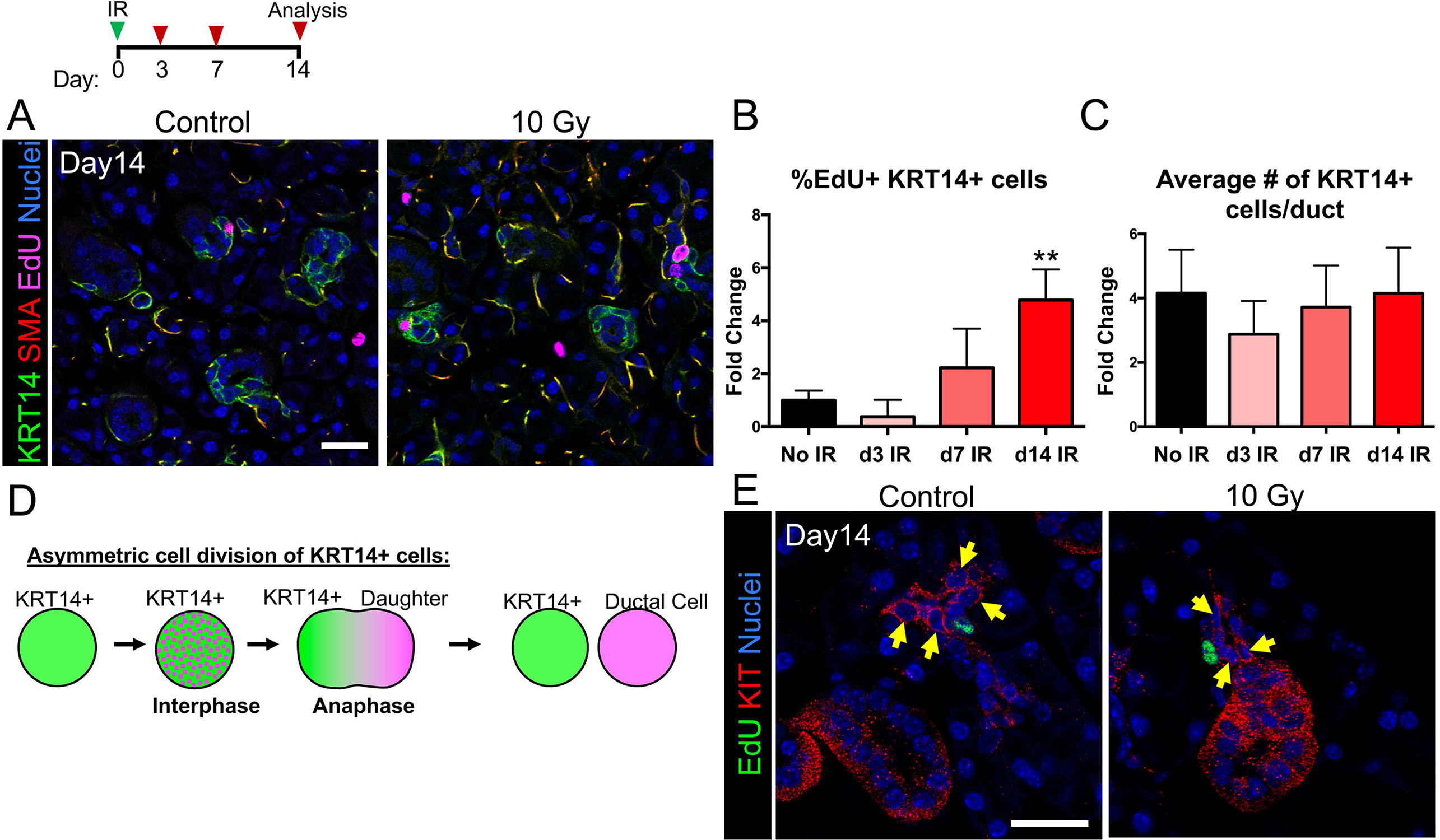
KRT14+ cells but not KIT+ cells proliferate after radiation and replenish the ductal compartment through asymmetric division. Adult C57Bl/6 mice were treated with 0 Gy (control) or 10Gy (IR) and sacrificed at 3, 7 or 14 days, with EdU being injected 2 h before collection. Number of proliferating cells (A) was then quantified (B and C). Data in B and C (n=3 mice) are means+s.d. and were analyzed using a one-way analysis of variance with a post-hoc Dunnett’s test. ** = p<0.005. (D) Proposed model of asymmetric division of KRT14+ cells. (E) No proliferation was found in KIT+ cells (yellow arrows) at any stage in control or irradiated SGs. Scale bar=25µm.

**Figure 7.**
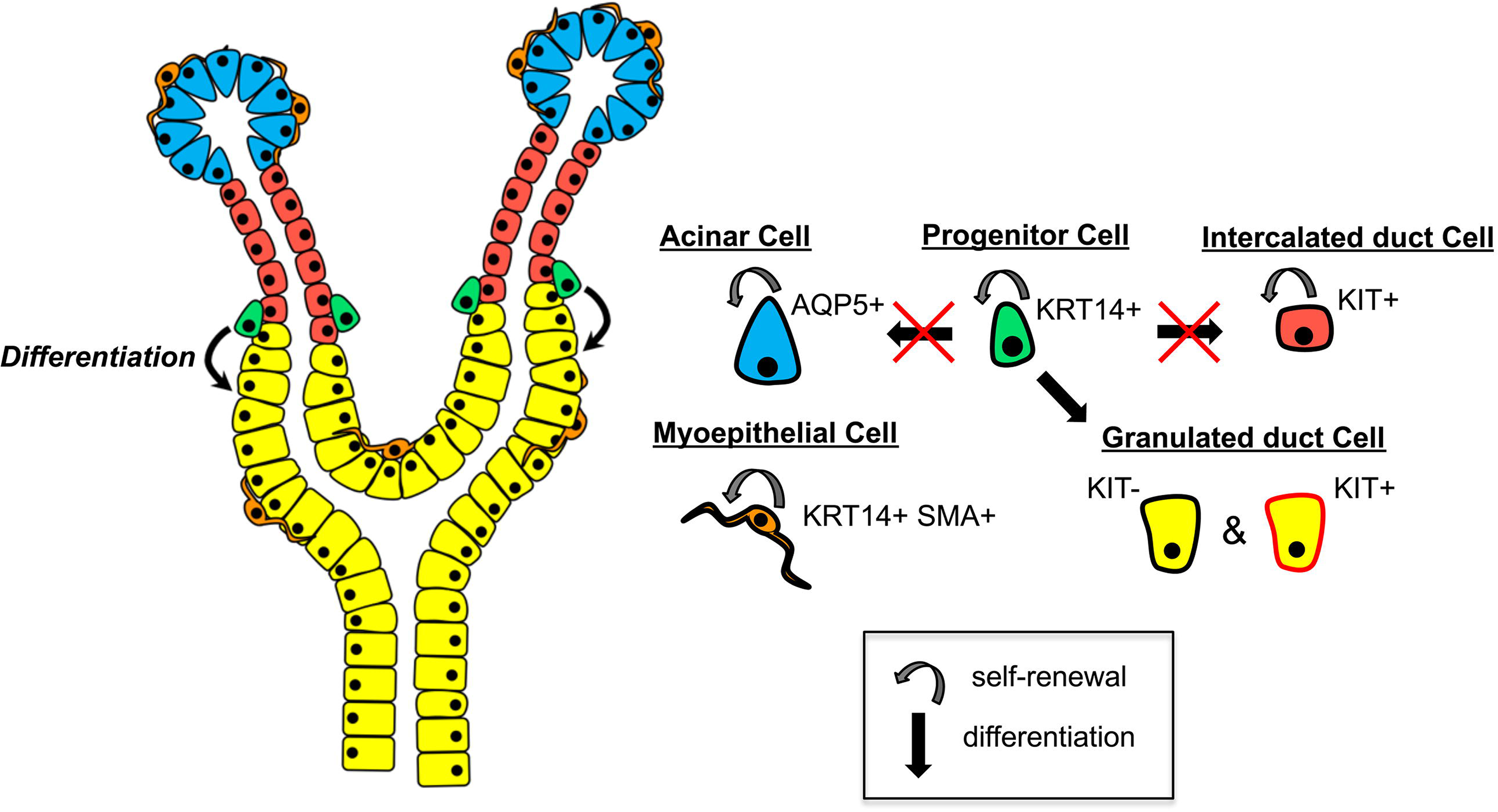
Schematic representation of the heterogeneous progenitor populations that maintain and replenish the adult SG under homeostatic and injury conditions. KRT14+ progenitor cells self-renew and give rise to differentiated KIT-negative and KIT-positive granulated duct cells but do not contribute to the acinar or intercalated duct (ID) compartments. KIT+ ID cells are long-lived and slow dividing, and maintain the ID compartment. KRT14+ SMA+ myoepithelial cells (MEC) self-renew and maintain the MEC population. AQP5+ acinar cells self-renew and replenish the acinar cell population.

## Discussion

A large number of studies over the last 70 years have suggested that the ductal system is comparatively more resistant to radiation-induced damage than the acinar cells (Redman 2008; Grundmann et al. 2010). Yet, whether this is the case had not been empirically determined. Our recent finding that acinar cells are capable of regenerating immediately after radiation exposure (Emmerson et al. 2018) strongly suggested that the ductal system could also regenerate in response to genotoxic injury. Consistent with this prediction, we show that ducts can indeed regenerate after radiation-induced damage, however, cell replacement primarily occurs in the granulated ducts and is mediated by KRT14+ cells. In contrast, KIT+ cells show little turnover to maintain themselves, possibly surviving through increased resistance to DNA damage mediated cell death to ensure tissue architecture remains unperturbed. Furthermore, we also find that these progenitors, like the SOX2+ acinar progenitors, do not gain lineage plasticity in response to radiation thereby ruling out a regenerative mechanism used by other organs such as the skin (Page et al. 2013; Ito et al. 2005) to ensure organ fidelity. Thus, these data indicate that the salivary ductal progenitor cells possess regenerative capacity but that there is heterogeneity in the mechanisms by which they maintain the organ.

Epithelial stem cells can divide by symmetrical or asymmetrical division, which allows for the expansion of their numbers and the production of differentiated cells (Itzkovitz et al. 2012; El-Hashash & Warburton 2012; Lechler & Fuchs 2005). An excellent example of this is the recent study of prostate basal and luminal cells: the basal cells exhibit both asymmetric and symmetric division to self-renew and produce differentiated luminal progeny whereas the luminal cells divide by symmetrical division to produce themselves (Wang et al. 2014). We have previously shown basal cells in the lacrimal gland undergo asymmetric division to produce luminal daughter cells (Farmer et al. 2017) and our data in the SG suggests a similar outcome for the basal KRT14+ cells lining the excretory ducts. Intriguingly, as the KRT14+ cells lie outside the granulated ducts, these daughter cells must incorporate into the larger ductal system and produce a cell vastly different in morphology from the original KRT14+ cell. Similar to the luminal cell of the prostate, our lineage tracing suggests KIT+ cells of the intercalated ducts undergo symmetric divisions (albeit very slowly) producing themselves over and over again albeit at a very slow rate. It is likely that symmetric division is slow in these cells due the confined size of the intercalated ducts which are typically only 10 cell lengths in size. Their long-lived nature (as well as progenitor cells status) is also supported by previous label retaining studies in adult rodents showing that intercalated duct cells retain BrdU for more than 7 weeks after their initial labeling (Chibly et al. 2014). Further investigation is necessary to understand why KIT+ and KRT14+ cells are restricted to symmetric and asymmetric divisions as well as the mechanisms regulating the quiescence of KIT+ cells.

We recently reported that murine acinar progenitor cells marked by SOX2 are highly regenerative, at least in the first 30 days after radiation exposure, and are capable of repopulating the acini similar to uninjured controls (Emmerson et al. 2018). Similarly, here we find that ductal system can also replenish itself to some extent after genotoxic shock through KRT14+ cells. KRT14+ cell proliferation increases in the 2 weeks after radiation exposure, suggesting there is feedback from the granulated ducts to promote replenishment of these cells. However, whether this regenerative capacity can be sustained for the long term is unclear. Previous studies indicating that murine salivary glands degenerate/senesce 3-6 months after radiation suggest ducts may only regenerate for a limited time (Marmary et al. 2016; Muhvic-Urek et al. 2006). As these investigations focused on the acini rather than ducts, it remains to be determined whether the regenerative capacity of KRT14+ cells does fail eventually and further analysis is required to discern the cause. It also remains to be determined whether the human salivary gland ductal system actively regenerates in the days/months after therapeutic radiation and if this regenerative capacity fails in the long-term due to the inability or progenitors to enter the cell cycle.

Our study indicates that genotoxic damage to the SG does not induce the acquisition of lineage plasticity, as has been observed after injury in the intestine (Van Es et al. 2012), trachea (Tata et al. 2013) and stomach (Stange et al. 2013), where cells repopulate the lost compartment irrespectively of their origin and function under unperturbed conditions. This presents a significant challenge for the tissue as lineage plasticity of functionally distinct stem cell populations is a robust fail-safe mechanism to maintain regenerative capacity in case of stem cell loss when tissue is damaged. However, it is also possible that salivary ductal progenitor cells possess a robust response to oxidative/DNA damage and are able to efficiently repair themselves to reduce this reliance on other mechanisms. Also, the close proximity of KIT+ to KRT14+ cells to each other suggests that they may behave as reciprocating niche cells, positively influencing the function of the other cell to indirectly promote repair. Further studies are required to understand their interactions and whether communication between these two cells is required for their homeostatic and regenerative capacity. In addition, whether these cells undergo plastic interconversion when under different injury conditions remains to be tested.

## Materials and Methods

### Mouse Lines

All procedures were approved by the UCSF Institutional Animal Care and Use Committee (IACUC) and were adherent to the NIH Guide for the Care and Use of Laboratory Animals. Mouse alleles used in this study were provided by The Jackson Laboratory and include *Krt14^CRERT2^* (Vasioukhin et al. 1999), *Kit^CreERT^* (Klein et al. 2013)*, Acta2^CreERT2^* (Wendling et al. 2009) and *Rosa26^mTmG^* (Muzumdar et al. 2007) *and Rosa26^RFP^* (Luche et al. 2007).

### Lineage Tracing of KRT14+ Cells

*Krt14^CreERT2^;Rosa26^mTmG^* embryos were generated by breeding *Krt14^CreERT^2*(Ki/+);mTmG(Ki/Ki) males with *Krt14^CreERT2^*(+/+);*Rosa26^mTmG^* (Ki/Ki) females. Timed pregnant females were injected with 5.0mg/20g tamoxifen (Sigma Aldrich) in corn oil (Sigma Aldrich) at E10.5 and E12.5 and euthanized at E16.5, or injected with tamoxifen at E15.5 or E16.5 and euthanized at E18. For postnatal lineage tracing of KRT14+ cells, P2 or P30 pups were injected with one single dose of 0.3mg or 5mg tamoxifen respectively and euthanized on P30 or P270. For adult lineage tracing of KRT14+ cells, *Krt14^CreERT2^*(Ki/+); *Rosa26^mTmG^* (Ki/Ki) females were injected with one single dose of 2.5mg/20g tamoxifen and euthanized after 24 hours or 14days.

### Lineage Tracing of KIT+ Cells

Postnatal lineage tracing of KIT+ cells was carried out by injecting P2 *Kit^CreERT2^*(Ki/+); *Rosa26^mTmG^* (Ki/Ki) pups with one single dose of 0.3mg tamoxifen and euthanizing on P30. For adult lineage tracing of KIT+ cells, *Kit^CreERT2^*(Ki/+); *Rosa26^mTmG^* (Ki/Ki) females were injected with 4mg/30g tamoxifen for three consecutive days and euthanized on day 14. For gamma irradiation studies, *Kit^CreERT2^*(Ki/+); *Rosa26^mTmG^* (Ki/Ki) adults were injected with one single dose of 2.5mg tamoxifen 24 hours before irradiation and euthanized 30 days later.

### Lineage Tracing of SMA+ Cells

*Acta2^CreERT2^;Rosa26^RFP^* embryos were generated by breeding *Acta2^CreERT2^*(Ki/+);*Rosa26^RFP^* (Ki/Ki) males with *Acta2^CreERT2^*(+/+);*Rosa26^RFP^*(Ki/Ki) females. Timed pregnant females were injected with 5.0mg tamoxifen at E15.5 and euthanized at E18. For adult lineage tracing of SMA+ cells, *Acta2^CreERT2^*(Ki/+); *Rosa26^RFP^* (Ki/Ki) males were injected with one single dose of 5mg tamoxifen and euthanized on 30 days or 6 months later.

### Gamma-Radiation

Gamma-radiation experiments of adult murine salivary glands was carried out as previously described (Emmerson et al. 2018). In brief, C57BL/6, *Krt14^CreERT2^*;*Rosa26^mTmG^* and *Krt14^CreERT2^;Rosa26^mTmG^* mice were anesthetized with 1.25% 2,2,2 tribromoethanol (Alfa Aesar) in 0.9% saline (Vedco Inc.) and irradiated using a ^137^Cs source in a Shepherd Mark I 68A ^137^Cs Irradiator (JL Shepherd & Associates). Only the neck and part of the head were exposed, and the salivary glands were radiated with two doses of 5 Gy at a dose rate of 167 Rads/min for 2.59 min (one of each side of the head, bilateral, and sequential but on the same day) for a total dose of 10 Gy. Control mice were anesthetized as per experimental mice, but did not undergo radiation treatment. All animals were allowed to completely recover before returning to normal housing and were given soft diet *ad libitum* (ClearH_2_O). Mice were euthanized after 3, 7, 14 and 30 days. For proliferation analysis, animals were i.p injected with 0.25mg/25g 5-ethynyl-2’-deoxyuridine (EdU – Thermofisher Scientific) 2 hours before sacrifice.

### Human salivary gland tissue collection

Human fetal salivary glands were collected from post-mortem fetuses at 22 weeks gestation with patient consent and permission from the ethical committee of the University of California San Francisco. Following dissection, salivary glands were fixed immediately in 4% PFA and fixed overnight at 4°C. Adult human salivary gland was obtained from discarded, non‐identifiable tissue with consent from patients undergoing neck resection (age 31yr old male). Informed consent was given by all subjects and experiments conformed to the principles set out in the WMA Declaration of Helsinki and the Department of Health and Human Services Belmont Report. Tissue was immediately transferred in ice cold PBS to the laboratory where it was fixed in 4% PFA.

### Tissue Processing

Embryonic SGs were dissected and fixed for 2 hours at room temperature, while adult murine, human fetal and adult human SGs were fixed overnight at 4°C, in 4% PFA. Tissue was incubated in increasing concentrations of sucrose (12.5%-25%) and embedded in in OCT (Tissue-Tek). Tissue was sectioned using a Leica cryostat and tissue sections kept at -20°C until immunofluorescent analysis. OCT tissue blocks were kept at -80°C.

### Immunofluorescent Analysis

Tissue sections were left to equilibrate at room temperature for 10 minutes and washed in PBS. Tissue sections were permeabilized in 0.5% Triton-X in PBS for 10 minutes and blocked for 2 hours at room temperature in 10% Donkey Serum (Jackson Laboratories, ME) and 1% BSA (Sigma Aldrich) in 0.05% PBS-Tween-20. For anti-c-KIT primary antibody staining, permeabilization with ice-cold 1:1 Acetone:Methanol solution was carried out for 1 minute, followed by incubation in blocking solution as above. Tissue was incubated overnight at 4°C in primary antibody in 0.05% PBS-Tween-20. The following primary antibodies were used: rabbit anti-SMA (1:200, Abcam, AB5694), mouse anti-SMA-Cy3 conjugated (1:400, Sigma Aldrich, C6198), rabbit anti-AQP5 (1:200, Millipore, AB3559), rat anti-E-cadherin (1:300, Life Technologies, 13–1900), rabbit anti-c-KIT (1:200, Cell Signaling, 3074), goat anti-c-KIT (1:200, Santa Cruz, sc-1494), rabbit anti-KRT5 (1:1000, Covance, PRB-160P), rat anti-KRT8 (1:400, Troma I, DSHB), rabbit anti-KRT14 (1:1000, Covance, PRB-155P), rat anti-KRT8 (1:200, Troma II, DSHB), rat anti-KRT19 (1:300, DSHB, troma III), goat anti-MUC19 (1:200, AbCore, AC21-2396). Antibodies were detected using Cy2-, Cy3- or Cy5-conjugated secondary Fab fragment antibodies (Jackson Laboratories) and DNA was labeled with Hoescht 33342 (1:1000, Sigma Aldrich). EdU staining was performed using the Click-iT EdU Alexa-Fluor 647 kit. Slides were mounted using Fluoromount-G (SouthernBiotech) and tissue was imaged using a Leica Sp5 confocal microscope and NIH ImageJ software.

### Cell Number and Proliferation Analysis

For cell number and proliferation analysis of KRT14+ and KIT+ cells, control and irradiated adult female tissue was stained using the Click-iT EdU kit (Thermofisher Scientific). Cells positively stained for EdU and cell markers were counted using NIH Image J software. All data was obtained from 3 fields of view from each animal, where 3 control and 3 irradiated mice were analyzed for each time point.

### Statistical Tests

Data were analyzed for statistical significance using a one‐way ANOVA (multiple groups) with *post hoc* testing performed using Dunnett (GraphPad Prism). For multiple testing, we used a false discovery rate of 0.05. All graphs show the mean + standard deviation (SD) and were generated using GraphPad Prism.

## Acknowledgements

The authors would like to acknowledge Eliza Gaylord, Sonia Sudiwala and Aaron Mattingly for their laboratory assistance.

## Author Contributions

AM and SMK designed and planned the study; AM, NPC, EE, SN and performed the experiments; AM and SMK analyzed and interpreted the data; KS and OK provided the SG from the *Acta2*^*CreERT2*^; *Rosa26*^*RFP*^ adult mice; and AM and SMK prepared the manuscript, with input from EE.

## References

Brown, L.R. et al., 1975. Effect of radiation-induced xerostomia on human oral microflora. Journal of dental research, 54(4), pp.740–750.

Chibly, A.M. et al., 2014. Label-retaining cells in the adult murine salivary glands possess characteristics of adult progenitor cells. J. A. Chiorini, ed. PloS one, 9(9), p.e107893.

Dreizen, S. et al., 1977. Prevention of xerostomia-related dental caries in irradiated cancer patients. Journal of dental research, 56(2), pp.99–104.

Dusek, M. et al., 1996. Masticatory function in patients with xerostomia. Gerodontology, 13(1), pp.3–8.

El-Hashash, A.H.K. & Warburton, D., 2012. Numb expression and asymmetric versus symmetric cell division in distal embryonic lung epithelium. The journal of histochemistry and cytochemistry : official journal of the Histochemistry Society, 60(9), pp.675–682.

Emmerson, E. et al., 2018. Salivary glands regenerate after radiation injury through SOX2-mediated secretory cell replacement. EMBO molecular medicine, p.e8051.

Farmer, D.T. et al., 2017. Defining epithelial cell dynamics and lineage relationships in the developing lacrimal gland. Development, 144(13), pp.2517–2528.

Grundmann, O. et al., 2010. Restoration of radiation therapy-induced salivary gland dysfunction in mice by post therapy IGF-1 administration. BMC cancer, 10(1), p.417.

Ito, M. et al., 2005. Stem cells in the hair follicle bulge contribute to wound repair but not to homeostasis of the epidermis. Nature Medicine, 11(12), pp.1351–1354.

Itzkovitz, S. et al., 2012. Optimality in the development of intestinal crypts. Cell, 148(3), pp.608–619.

Kim, Y.-J. et al., 2008. Comparative analysis of ABCG2-expressing and label-retaining cells in mouse submandibular gland. Cell and Tissue Research, 334(1), pp.47–53.

Kimoto, M. et al., 2007. Label-retaining Cells in the Rat Submandibular Gland. The journal of histochemistry and cytochemistry : official journal of the Histochemistry Society, 56(1), pp.15–24.

Klein, S. et al., 2013. Interstitial cells of Cajal integrate excitatory and inhibitory neurotransmission with intestinal slow-wave activity. Nature communications, 4, p.1630.

Knox, S.M. et al., 2010. Parasympathetic innervation maintains epithelial progenitor cells during salivary organogenesis. Science (New York, N.Y.), 329(5999), pp.1645–1647.

Knox, S.M. et al., 2013. Parasympathetic stimulation improves epithelial organ regeneration. Nature communications, 4, p.1494.

Kwak, M., Alston, N. & Ghazizadeh, S., 2016. Identification of Stem Cells in the Secretory Complex of Salivary Glands. Journal of dental research, 95(7), pp.776–783.

Lechler, T. & Fuchs, E., 2005. Asymmetric cell divisions promote stratification and differentiation of mammalian skin. Nature, 437(7056), pp.275–280.

Lee, M.G. et al., 2012. Molecular mechanism of pancreatic and salivary gland fluid and HCO3 secretion. Physiological reviews, 92(1), pp.39–74.

Lombaert, I.M.A. et al., 2013. Combined KIT and FGFR2b signaling regulates epithelial progenitor expansion during organogenesis. Stem cell reports, 1(6), pp.604–619.

Luche, H. et al., 2007. Faithful activation of an extra-bright red fluorescent protein in “knock-in” Cre-reporter mice ideally suited for lineage tracing studies. European Journal of Immunology, 37(1), pp.43–53.

Marmary, Y. et al., 2016. Radiation-Induced Loss of Salivary Gland Function Is Driven by Cellular Senescence and Prevented by IL6 Modulation. Cancer research, 76(5), pp.1170–1180.

Muhvic-Urek, M. et al., 2006. Imbalance between apoptosis and proliferation causes late radiation damage of salivary gland in mouse. Physiological research, 55(1), pp.89–95.

Muzumdar, M.D. et al., 2007. A global double-fluorescent Cre reporter mouse. Genesis (New York, N.Y. : 2000), 45(9), pp.593–605.

Page, M.E. et al., 2013. The epidermis comprises autonomous compartments maintained by distinct stem cell populations. Cell stem cell, 13(4), pp.471–482.

Redman, R.S., 2008. On approaches to the functional restoration of salivary glands damaged by radiation therapy for head and neck cancer, with a review of related aspects of salivary gland morphology and development. Biotechnic & histochemistry : official publication of the Biological Stain Commission, 83(3-4), pp.103–130.

Siegel, R.L., Miller, K.D. & Jemal, A., 2015. Cancer statistics, 2015. CA: A Cancer Journal for Clinicians, 65(1), pp.5–29.

Srinivasan, R. & Chang, W.W.L., 1979. The postnatal development of the submandibular gland of the mouse. Cell and Tissue Research, 198(2).

Stange, D.E. et al., 2013. Differentiated Troy+ chief cells act as reserve stem cells to generate all lineages of the stomach epithelium. Cell, 155(2), pp.357–368.

Sullivan, C.A. et al., 2005. Chemoradiation-induced cell loss in human submandibular glands. The Laryngoscope, 115(6), pp.958–964.

Tata, P.R. et al., 2013. Dedifferentiation of committed epithelial cells into stem cells in vivo. Nature, 503(7475), pp.218–223.

Tucker, A.S., 2007. Salivary gland development. Seminars in Cell & Developmental Biology, 18(2), pp.237–244.

Van Es, J.H. et al., 2012. Dll1+ secretory progenitor cells revert to stem cells upon crypt damage. Nature cell biology, 14(10), pp.1099–1104.

Vasioukhin, V. et al., 1999. The magical touch: genome targeting in epidermal stem cells induced by tamoxifen application to mouse skin. Proceedings of the National Academy of Sciences of the United States of America, 96(15), pp.8551–8556.

Wang, J. et al., 2014. Symmetrical and asymmetrical division analysis provides evidence for a hierarchy of prostate epithelial cell lineages. Nature communications, 5, p.4758.

Wendling, O. et al., 2009. Efficient temporally-controlled targeted mutagenesis in smooth muscle cells of the adult mouse. Genesis (New York, N.Y.: 2000), 47(1), pp.14–18.

